# Examining Academic Leader’s work in implementing Competency-based Medical Education using Organizational Learning Theory

**DOI:** 10.1101/438077

**Authors:** Nicolas Fernandez, Nicole Leduc, Nathalie Caire Fon, Louis-Georges Ste-Marie, Dat Nguyen-Dinh, Andrée Boucher

**Affiliations:** Dr. Fernandez is Assistant Professor at the Center for Pedagogy Applied to the Health Sciences and at the Department of Family Medicine and Emergency Medicine of the Faculty of Medicine, at the Université de Montreal, Montréal, Canada.; Dr. Leduc Nicole, is Full Professor in the Department of Health Management, Evaluation and Policy, at the School of Public Health, at the Université de Montreal, Montréal, Canada.; Dr. Caire Fon, Nathalie, is Adjunct Professor in the Department of Family Medicine and Emergency Medicine, at the Faculty of Medicine, at the Université de Montreal, Montréal, Canada.; Dr Ste-Marie is Full Professor in the Department of Internal Medicine at the Faculty of Medicine, at the Université de Montreal, Montréal, Canada; Dr Nguyen-Dinh, Dat is a Family Physician in Montreal, Québec, Canada.; Dr Boucher is Full Professor in the Department of Internal Medicine at the Faculty of Medicine, at the Université de Montreal, Montréal, Canada.

## Abstract

**Context:** Competency-based medical education (CBME) implementation is being carried out in many medical schools worldwide. Academic Leadership is a strategy where selected Faculty act to influence peers to adopt change. The Université de Montréal medical school, has adopted this strategy to implement CBME.

**Purpose:** This paper aims to describe the work of Academic Leaders in the process of CBME implementation and to explore relevance of the Nonaka and Toyama organizational learning theory to map implementation progress.

**Method:** Because knowledge creation model focuses on the relationships between leaders and social structures, embedded case study was selected. Diverse sampling method was used to select three departments: internal medicine, surgery and psychiatry, based on the number of CBME training activities. Data collection was at two intervals, two years apart. Semi-structured interviews (individual and group) were conducted with Department Heads and Academic Leaders. Thematic analysis was conducted on the 15 interview transcriptions.

**Results:** As implementation begins, Leaders critically revisit accepted teaching routines and develop a common conception of CBME. This enables leaders to communicate with a wider audience and work within existing committees and working groups where they “break down” CBME into practical concepts. This practical understanding, disseminated through Entrustable Professional Activities, enables observable change.

**Conclusion:** Leaders’ roles evolved from an “expert” that disseminates knowledge about CBME through lectures, to a responsive and pragmatic supporting role by developing and writing practical tools in collaboration with peers and program directors.

## Introduction

One the difficulties medical schools have encountered in implementing Competency-based medical education (CBME) stems from the fact that CBME frameworks describe wide scoping curricular outcomes with little guidance to instructors on the ground on how to teach and assess competencies (1–6). This lack of practical guidance generates much resistance towards CBME implementation (7) and little attention has been paid in the scientific literature to organisational strategies employed to attenuate the resistance to change. This study describes how Academic Leadership can bridge the gap between CBME concepts and practical tools that instructors can use on the ground.

### Leadership in Higher Education

Studies in the field of leadership in higher education suggest that leadership in academic organizations is shifting towards a cultural perspective, based predominantly on relationships (8): ‘Without doubt, the key to being an effective leader is an ability to connect with others in the organisation and gain their cooperation in working collaboratively towards the organisational goals and objectives’ (p.29). Another perspective, that of transformational leadership, suggests that leaders communicate a vision, provide support, empower others to innovate and lead by example (9).

Among the strategies relied upon to support innovation in Higher Education (10) and CBME implementation in medical schools is one that rests on Grassroots Leaders (11) or Academic Leaders (12, 13). This approach consists of training a select group of faculty, generally interested and motivated by the vision of change promoted within the institution. The training can take the form of formal lectures on topics, group mentoring with an Educational Expert, or even graduate studies in medical education. Although Academic Leaders are generally not in a position of authority over their peers (14), they are expected to influence, motivate and inspire colleagues in adopting and successfully implementing change within a university department (15–17). This study proposes a theory guided exploration of the work of such a group of Academic Leaders as they work towards CBME implementation in a large medical school.

### Organizational change as knowledge creation

In our study, CBME implementation is construed as organizational change (18) within the medical school. We turned to the organizational development field that seeks to uncover levers for change (19, 20) provide a systematic perspective on the process. Some organizational change frameworks focus on power relationships and structures within the organization (21), others on the organization’s identity and interactions with the surrounding environment (22, 23) and still others on communities of practice (24, 25) and developing new knowledge (26). In contrast to the former, this latter perspective construes the individuals in an organization as knowledge creators.

A medical school’s decision to implement CBME does not come with clear definitions of how teachers are to conduct their teaching nor how they are to assess students’ competencies. New knowledge, generated from testing new conceptions of teaching and assessing, is required to solve the problem. As medical instructors experiment, exchange and discuss their insights with their colleagues, they embark on the process of organizational knowledge creation.

Tacit knowledge (27, 28), which hinges on the idea that an individual knows more than they can tell, is a key concept. Individuals develop personal insights into unique ways to conducting their work, which is reflected in the dual consciousness that characterizes humans at work: A practical consciousness, embedded in tacit knowledge, which informs what they do and a discursive consciousness that informs what they say (21) that is shared with others. Knowledge creation is conceptualized as the process of narrowing the gap between practical and discursive consciousness, or making tacit knowledge explicit. Unless there are overt efforts to facilitate this process, individuals’ tacit knowledge remains an untapped resource.

Hence, the challenge in implementing CBME is making the tacit knowledge held by instructors explicit and available to colleagues within the organization. This process can be referred to as organizational knowledge creation which rests on the idea ‘that knowledge is created and expanded through social interaction’ (29)(p. 61). This process, also referred to by Nonaka and colleagues as knowledge conversion, cannot be understood as a linear process, but as a spiral that expands organizational knowledge and distributes from individual, to group, to organisation and interorganisational levels (30).

### The knowledge creation process

It is implied that knowledge creation is not an individual activity and that it occurs within a given space and time, what Nonaka calls *ba*(30). *Ba* is a shared mental space and time where information is shared and interpreted collectively. The knowledge creation process is comprised of four stages that unfold within the *ba*: socialization, externalization, combination and internalization (SECI) which reflect a cumulative upward spiralling process.

The socialization stage emerges through day-to-day experiences in teaching medical students in a clinical setting. For example, medical educators become aware of contradictions in the way teaching is carried out and talked about. Talking to peers brings these contradictions to the fore. The socialization stage ends when individuals embrace actions to ‘resolve these contradictions’ (31)(p. 4).

During the externalization stage, individuals use their discursive consciousness to rationalize and articulate the contradictions they have encountered. Nonaka and Toyama insist that at this stage, individuals ‘seek to detach themselves from routines’ (p.4) to eventually put new ones in their place. Practical knowledge is actively shared with peers in an effort to form new concepts. The externalization stage ends when these new concepts becomes clear enough to be practical.

In the combination stage concepts are tested and disseminated to other members of the organization. This stage is where leaders ‘break down’ concepts, such as CBME, so that peers can develop their understanding of them in their discursive consciousness and make sense of them. The combination stage ends when peers take ownership of the new concepts and introduce them in their practice.

In the internalization stage newly created concepts generate new practical knowledge. The change in the organization becomes visible as a greater number of members adopt change and new routines take hold. This stage ends when individuals begin to critically question theses routines and a new cycle of knowledge creation begins. Table 1 summarizes the four stages of the knowledge creation model.

**Table 1:** The Four Stages of the Knowledge Creation Model (read clockwise)

| 1. Socialization | 2. Externalization |
| --- | --- |
| Sharing practical knowledge through direct experience (practical consciousness). | Articulating practical knowledge through dialogue and reflection (discursive consciousness). |

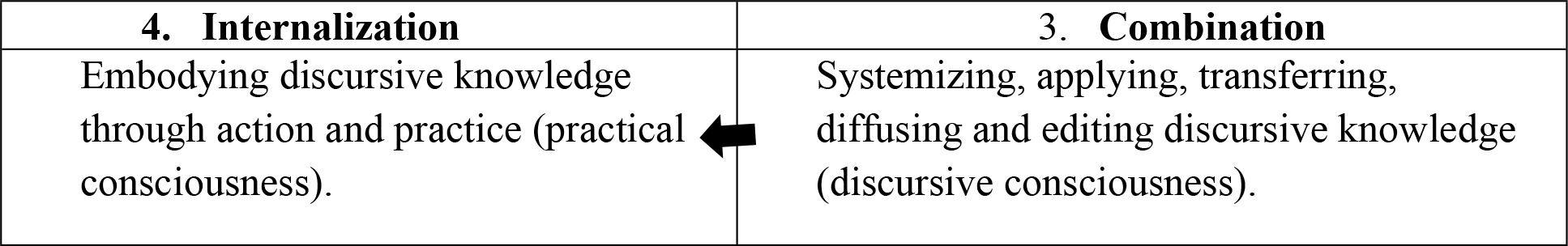

To summarize, an organization creates knowledge through the knowledge conversion process (SECI) which occurs in a specific time and space (*ba)* in the interactions between individuals and groups.

### Study aim

Some institutions of Higher Education make use of Academic Leaders to catalyse progress towards organisational change (32, 33). Yet, we know little about the ways in which Academic Leaders in universities develop and sustain the conditions and processes that generate change.

In our study, Academic Leaders, drawn from medical school faculty, were trained in CBME and were expected to guide and support their peers. How do they articulate their ideas of change and ultimately persuade peers to adhere to them?

Our study aimed to describe Academic Leaders’ work through the organisational knowledge creation process : socialization, externalization, combination and internalization. We were interested in identifying factors that enabled or hindered the transition from one stage of the process to the next. This would yield deeper insight into how Academic Leaders contribute to enacting change in higher education institutions.

## Methods

The largest francophone medical school in North America has undertaken CBME implementation and has chosen a strategy based on Academic Leaders. Academic Leaders’ mandate is to facilitate CBME implementation by supporting peers, developing and delivering training activities and advising Program Directors. Faculty of the medical school is made up of approximately 3000 instructors (roughly 20% tenured, 80% part-time or non-tenured), working in over 100 different teaching sites (hospitals, family medicine and community clinics, etc.) and offering 73 programs, from the undergraduate MD program to 72 graduate (residency and specialty) programs. The Faculty comprises 16 academic departments as well as two health sciences schools. Leaders who step-up from the ranks were trained in CBME and offered remuneration for their work.

In order to achieve the aims of the study, a case study approach with embedded levels (34, 35) was selected. The unit of enquiry is thus not the individual leaders, but the academic department to which they belong. This approach allowed observation of the interactions between the leaders and the social structures in which they work, which is a crucial dimension to the organizational knowledge creation model. We focused on leaders’ interactions with Department Chairs and with colleagues.

Departments were selected for intense study using a purposive sampling method (35)(p. 88). The small number of departments (N=16) makes random sampling unreliable because there is no way to ensure that these cases are representative of a larger population. In order to meet the goals of reproducing the relevant causal features assumed to be in the larger population and providing sufficient variation along the dimension of interest (CBME implementation), the diverse case sampling method was used (35)(p. 97). Hence, the departments were selected on the basis of a range of levels of CBME training activities in 2012 roughly equivalent to high, medium and low (35)(p. 98). The selected departments were medicine, surgery and psychiatry. The sample comprised a department with the highest number of training activities (MED), one that held little activity, (SURG) and one that had no training activities at all (PSY) in 2012.

The three Department Chairs were senior physicians (cardiologist, urologist and psychiatrist). All three had been involved in medical education, as instructors, program directors and Department Chairs for more than 10 years. When CBME implementation was announced in 2010, all three were already Department Chairs.

The 12 Academic Leaders interviewed were mid-career medical educators who demonstrated an interest for pedagogy. Two of the Academic Leaders had a graduate degree in medical education, the rest had gone through all the relevant CBME training sessions. One of them had been program director for many years and regularly facilitated training sessions on education for first year residents.

**Table 2:**
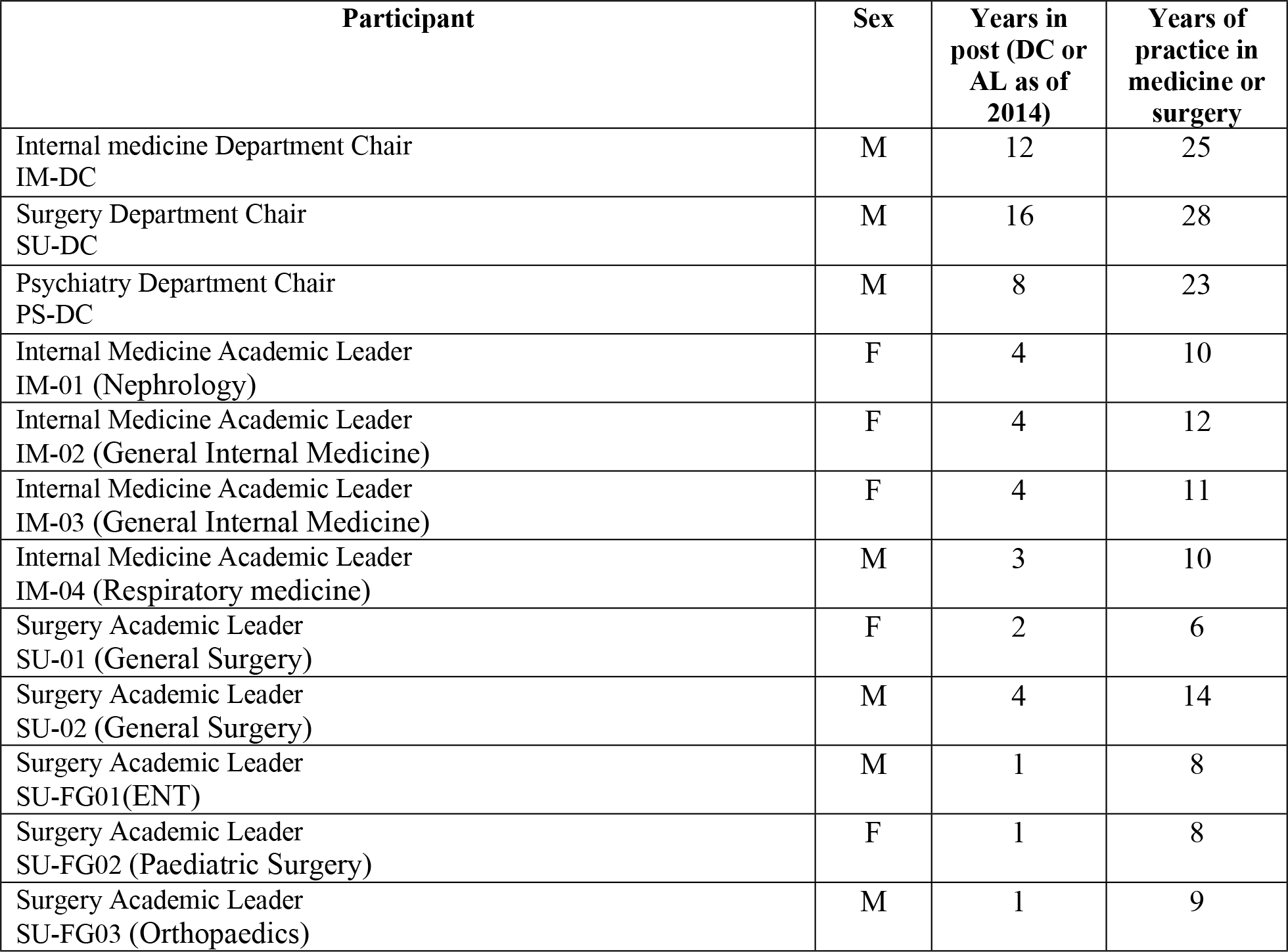
Study Participants

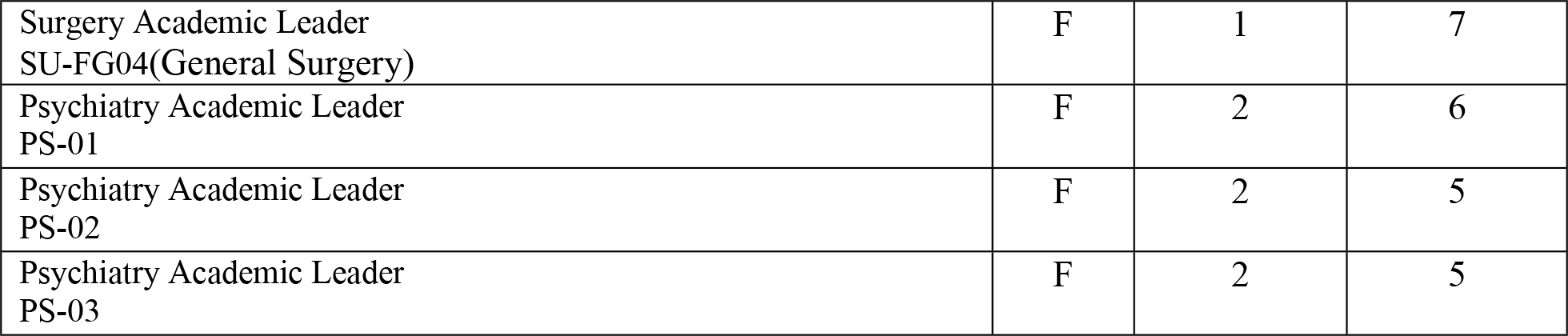

### Data collection and analysis

In order to apprehend the evolution of the work of Academic Leaders in CBME implementation over the two-year period of the study, two waves of data collection were carried-out. All three Department Chairs were interviewed in 2012 and in 2014. Three leaders of the internal medicine department were interviewed individually in 2012 and in a group interview in 2014. Two academic leaders in the surgery department were interviewed individually in 2014 and four different leaders were interviewed as a group in the same year. Two leaders of the psychiatry department were interviewed individually in 2014. In total, 13 semi-structured individual and 2 group interviews yielded the qualitative data that was recorded and transcribed. Questions and prompts focused on their experience and the facilitators and barriers they faced as they worked to implement CBME. All interviews where held on the main campus of the university.

**Table 3:**
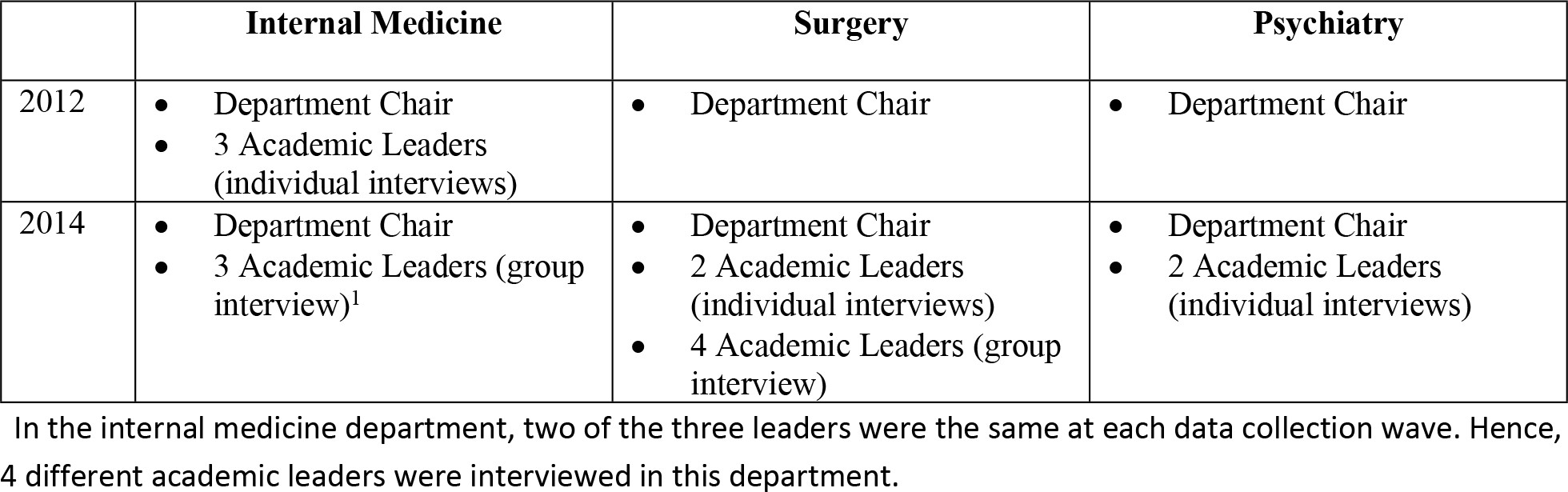
Data Collection Strategy

Thematic analysis was selected to code and identify common themes embedded in the transcripts of all 15 interviews (36, 37). Analysis was performed in three steps. In step one (initial review), one member of the research team (NF) identified and described themes that emerged from the data set. To limit the effect of our own biases, in step two, all themes were examined independently by two other members of the research team (NL, DN) to label and categorize each extraction until theme saturation was achieved. To complete step two, the research team reviewed the independently created themes and discussed their interpretation according to the research questions. Disagreements were resolved through discussions until a consensus was achieved. In step three, all codes were reviewed for accuracy and consistency with regards to the themes that emerged.

Ethical approval was sought and granted from the University Research Ethics Board (12-079-CERES-D).

## Results

In the following section we present the results of our analysis of the interview data of 2012 and 2014, revealing a more in-depth view of the work of Academic Leaders within the four knowledge conversion stages.

### CBME Implementation in 2012

Since 2005, school administrators had been announcing that CBME would be implemented and that funds were being set aside for the Academic Leaders in order to achieve this. In 2010, some Academic Leaders signed contracts with their Department Chair and the Vice-Dean for Continuing Medical Education (CME) that stipulated that they were to assist in Faculty development by supporting CBME implementation.

In 2012, only Academic Leaders that belonged in the Internal Medicine Department were active. In the other Departments, there had been some false starts. For example, Academic Leaders in the Psychiatry Department had signed contracts, prepared and delivered lectures on topics that they considered interesting. The reception was lukewarm at best. By 2012, they had given up and turned to other projects. Within the Surgery Department, Academic Leaders were just starting to be recruited in 2012.

In, 2012, the Department Chairs, recognized that the sheer number of medical instructors (approximately 3000) and the geographical distribution of the teaching sites represented in itself a major obstacle for the Leaders. This meant that, in order for CBME to be implemented, Leaders needed to reach out to colleagues within their own Department that were geographically spread-out:

> ‘If we want to reach out to instructors of the residents in any specialty, we have to have an impact across the board, throughout all [teaching] hospitals’. (SU-DC/2012)

Department Chairs, who allocated part of the development budget to compensate Academic Leaders, saw value in having them in their department and recognized that Faculty appreciated their presence. But, they felt that there was not enough of them to have a real impact. How can we ‘be serene enough to think of improving teaching when we don’t even have enough [resources] to write a report.’ (PS-DC/2012) At this initial stage, in the minds of Department Chairs, the lack of financial resources were considered an obstacle to CBME implementation.

Department Chairs mentioned changes in the vocabulary that now included many words and ideas associated with CBME. They noted a renewed interest in teaching among Faculty, but their enthusiasm ended there:

> ‘I don’t think those competencies are necessarily […] new, it’s perhaps another way of thinking, we agree, but we’ve known for years what a good doctor does. […] We knew what he is capable of doing with his patients, that he is capable of…. We knew it, it’s just [now it’s] expressed in a new framework…’ (IM-DC/2012)

Department Chairs were not quite certain whether CBME implementation was necessary at all. They repeatedly revealed the opinion that CBME is here today and gone tomorrow:

> ‘It’s not worth the investment to improve teaching. When we finish introducing the changes, we’ll have to start all over again because something new will come out. For those who are passionate about it, teaching is always evolving, constantly challenging assumptions. This was so even before we heard about Competency-based medical education.’ (SU-DC/2012)

Department Chairs regretted the absence of clear metrics to assess the impact of Academic Leaders’ work. They recognized that a training session given by them could attract many Faculty members, but the participation rates observed at the time weren’t high enough to fully justify the investment. Leaders themselves realized that the same people, interested in CBME, were recurring participants in their lectures. There still was a substantial number of Faculty who conveniently avoided lectures on the topic.

This rather pragmatic outlook contrasted sharply with the dynamic view held by Academic Leaders in 2012. We have to underline that the three internal medicine Leaders interviewed in 2012 were already convinced of the relevance of CBME. Their engagement in the program reflected a desire to improve teaching practice and manifested itself in the assumptions they challenged and the contradictions they singled out.

This initial interest was seen as crucial, but other elements emerged that facilitated or hindered Academic Leader’s involvement. Training on CBME provided by the Center for Pedagogy Applied to the Health Sciences (CPAHS) was adequate. CPAHS also provided logistical support for organizing events, providing learning materials and arranging for accreditation as Continuing Medical Education. Leaders recognized that without this support, their work would have been much more difficult:

> ‘The CPAHS is neutral and can provide some legitimacy when paired with a leader’ (PS-02/2014).

Furthermore, Leaders recognized that they had the support of their superiors and colleagues in their own teaching site. They felt that the organization valued pedagogy, and appreciated the specialized knowledge that they brought.

At this early stage, Academic Leaders didn’t expect to play an advisory role. They said that colleagues came to see them when they had questions. However, they didn’t feel at all prepared to play the expert role:

> ‘I barely have a grasp of competency-based concepts myself, I don’t see how I can sit with a program director and tell him what to do.’ (IM-01/2012)

As such, in 2012, Leaders started to train colleagues in their immediate teaching site. Getting the message out to colleagues in other teaching hospitals was definitely deemed harder than talking to colleagues in the same work place. ‘It was really hard to sit with them and ‘push’ CBME.’ (IM-01/2012)

Generally, people responsible for teaching in the hospitals called for Academic Leaders to give lectures. The lectures, Leaders admitted, were aimed at ‘bluntly exhorting teachers to change their teaching habits’ (IM-03/2012). At first they simply showed up ‘with [their] powerpoint, gave their talk, and left.’ (IM-01/2012). This generated much resistance, especially amongst physicians from other specialties. ‘Doctors don’t like to be told what to do, especially by someone who’s not from their specialty.’ (IM-02/2012)

Hence, in 2012, the bulk of Leaders’ activities were lectures, designed by them and aimed at ‘informing’ their colleagues about CBME. The issues they tackled stemmed from their representations of what CBME should be and were very often ‘interesting, but hardly concrete for what Faculty was supposed to be doing.’ (PS-DC/2012) Finally, Department Chairs had a sombre view of how Academic Leaders were going to achieve CBME implementation.

### CBME implementation two years later

In 2014, Academic Leaders were active in all three departments. In the following section, we present the results of the second set of interviews, conducted between March and April of 2014.

#### The Internal Medicine Department

As presented in the precious section, the socialization stage in the Internal Medicine Department was well underway in 2012. Active Academic Leaders had identified the reasons they wanted to get involved and they had started to identify a common vocabulary and goal.

Within the Internal Medicine Department, Academic Leaders, were seen as resource persons that you can go to if you have a problem. Academic Leaders acknowledged that colleagues were open to improving teaching and assessment:

> ‘We are identified as people who completed the training [on CBME], who give the training, people come to us for help. (…) I don’t know what changed, but there is an evolution where people are more motivated to do better, give students their feedback [on competencies]. ‘(IM-FG/2014, 143-147).

When the Department coordinated the effort to implement Entrustable Professional Activities (EPAs) (38, 39) it became clear that the combination stage had begun. EPAs, developed by the office of the Vice-Dean for Continuing Medical Education and the CPAHS, were formative assessment tools that enabled transposition of the abstract competencies into teachable and assessable moments. Academic Leaders were very active in developing and implementing EPAs in many internal medicine programs and CBME was becoming a ‘hot’ topic in Continuing Medical Education activities.

Another indication of the combination stage is the recognition that Academic Leaders received from within their Department. They became known as people who know about CBME ‘people willing to help, willing to motivate people.’ (IM-FG/2014). Even residents (graduate students) come to Academic Leaders when they have a problem with teaching. They come to see them because they know they’ve been working on CBME but not because they are Academic Leaders.

> ‘Me, people come to see me because they see me everywhere and they know that I’ll take care of it if they come and see me and they feel comfortable to talk about it with me. But I don’t think that the “Academic Leader” tag has changed anything.’ (IM-FG/2014)

By 2014, Internal Medicine Academic Leaders increasingly positioned themselves in coordinating positions, but remained connected to what was happening in teaching sites as they figured out what it took to reduce colleagues’ resistance: ‘Personally, I think there must be local initiatives with people that are well acquainted with local issues and this is how we get professors interested.’ (IM-FG/2014) They had clearly passed the threshold into the combination stage as they were having a wider impact within the medical school thanks to the support they provided in writing EPAs.

#### The Psychiatry Department

In 2012, Academic Leaders in the Psychiatry Department had stopped working and their contracts had not been renewed. Two years later, three different instructors got involved because they were interested in pedagogy from the outset. One of them stated that pedagogy is ‘a stimulating and interesting field that touches everybody.’ (PS-01/2014). As another put it, CBME implementation triggered discussions about pedagogy among colleagues in her teaching hospital, ‘it was like venting the ground that hadn’t been turned in 20 years.’ (PS02/2014)

Leaders also made it clear that their involvement didn’t depend exclusively on the availability of financial resources. Pedagogy was considered sufficiently important that they would get involved even without contracts or the Academic Leader title. This is reflected in the following quote about a well-known fact in academia: ‘When we submit a funding proposal for research, even if we don’t get the funding, we still do the research’ (PS-02/2014).

One of the Academic Leaders brought up issues of legitimacy associated with the role of ‘expert’. This created a distance from her colleagues:

> ‘we have to find a way to define this institutionally […] we are quickly placed in the position of the one who knows. This is the way things happen at the university, you go and do a presentation on anxiety troubles, even if your purpose is to do prevention and not…well, suddenly you’re the expert.’(PS-02/2014)

By 2014, the three Academic Leaders realized they needed to work together in association with the Department Chair. Consequently, they progressed through the socialization stage from an intuitive approach, seeing leadership as an individual endeavour, to a shared leadership approach better suited to bring about change in a large academic department.

With the help of the Department Chair, Academic Leaders in psychiatry had managed to get liberated time. This allowed Academic Leaders to reach beyond their department and ask the CPAHS for assistance. They were provided with ideas and tools, which had a considerable impact on their work

> ‘Very often we speak of…. When I made a presentation in my hospital, it was very local, at first. I didn’t have a clear goal….I just had the impression that it was a need everywhere, and so, I said: ‘Oh, finally!’ In actuality, we work with many of the things that are provided us by the CPAHS’ (PS-01/2014).

They were also involved in larger initiatives for CBME implementation such as defining the competency framework and participated in meetings with colleagues from other departments. They had also introduced themselves to the academic committees at the psychiatry teaching hospitals:

> ‘It was really to train people on competencies in all programs so that there could be people who piloted the project somewhat….but I think that at that moment there wasn’t a clear vision yet. In any case, that’s the impression I was under. We had a project but it hadn’t yet taken shape. So, we were three [of us] at the time and what we did is that we went round to all teaching sites to present competencies to our colleagues. And, what we received in terms of feed-back, already […] psychiatrists were interested, there was some resistance: again, something new that we had to teach, competencies and all. But there was an interest.’ (PS-01/2014).

Acknowledgment from peers was an important issue for Leaders in this department, especially since they had to intervene in settings that were not their own. There was a sense that leaders were sandwiched between the Department Chair and the colleagues on the ground, they felt this took away some traction.

> ‘…it’s never simple to be recognized as a leader when we were designated by someone else (…) at the same time I had the support from the Department Chair and…of the Vice-dean, but at the same time work on the ground for implementation of the concept of leader even…of…implementing pedagogical instructions was just in its infancy, it was just beginning, so that at the same time we have a role, but we weren’t sure where we stood.’ (PS-02/2014)

Working as a group acted as a catalyst. It provided the structure and legitimacy required to push for changes and members became recognizable to colleagues as people that could help with CBME. It was clear to the leaders that coordinating team activities was crucial ‘to figure out what we had to do’. (PS-02/2014) They pointed out that working as a team avoids getting dispersed and waste energy. If we work on our own, we get easily ‘distracted by our regular duties.’ (PS-01/2014) Leaders clearly felt that their decision to work together was an important facilitating factor.

> ‘what helps, is not to be alone. The group’s impulse and the fact that we are a few of us to carry this mandate…because with such a big department, so widespread, at one point we don’t know where to start.’ (PS-01/2014)

The decisive moment came when the Academic Leaders in the psychiatry department realized that faculty development and Continuing Medical Education (CME) should go hand in hand.

> ‘It was natural for us to join aspects of Academic Leadership and Continuing Medical Education (CME), and we said: let’s have a common vision! […] I decided to look at CME from the perspective of pedagogy. (…) I said that for me CME will be like Faculty development. Ok, this suited me. So, I would say that it was in 2012 that things started-up again. And in 2013 we had a pretty productive year thanks to this renewed energy that we felt. And whether you want it or not, we were under pressure….we felt that psychiatry was lagging behind, we really felt it…’(PS-01/2014)

By 2014, Academic Leaders in the psychiatry department had set up working committees in all programs of the department. It became clear to them that they needed to partner with teaching sites and the instructors within them. ‘The fact that we had representatives in the teaching hospitals that were engaged also kept us going.’ (PS-01/2014)

Because of the number of teaching sites, it was impossible to think of embedding a leader in each site. Leaders understood the necessity to partner with teachers on the ground.

> ‘So, this is when we finally, [realized] it’s just us three. We sometimes got time off and worked some files a little more, then we brought it back to a wider group because we thought that it was important that the work be done at each teaching site. […] We worked more in detail […] and we submitted our work to the committee and the committee made comments, and we came back. […] and so it was that last autumn we organized a whole day for the implementation of EPAs […] we managed to get about 50 psychiatrists who answered the call to whom we presented the EPAs and started working on the generic EPAs, chose the ones that were most relevant for [psychiatry] and got all our groups working. We have working groups in psychiatry who are responsible for the mandatory rotations […] and all rotations were represented by the sub-committee and in those sub-committees, they appropriated themselves the EPAs. (PS-01/2014, 183-194)

Having introduced themselves on all program committees, Academic Leaders were actively collaborating in EPA development, providing guidance and support:

> ‘I told the Vice-Dean that I appreciated the extraprofessional exchanges […] this should be included in our teacher training, it could be something that could […] enrich what we try to give because there is what we can present and lecture about concerning ways of teaching, but what people talk about amongst themselves, that’s even more enriching.’ (PS-01/2014)

Here Academic Leaders signal a change in the way they work, based on listening to what colleagues have to say far more than trying to exhort them to change. Leaders expressed that the key was being present in committees, whether they be university committees or hospital committees. ‘The aim is to get people thinking of pedagogy.’ (PS-02/2014)

As CBME implementation progressed, Academic Leaders felt that they were making a difference. At the very least they were under the impression that CBME is getting simpler and concrete. They realized that what works best is to bring concrete worked-out solutions and make colleagues feel that what you are proposing is going to formalize what they are already doing. This can be associated with the transition from the externalisation to the combination stage, where concepts are disseminated to other members of the organization who gain a practical understanding of them in their discursive consciousness. Leaders felt that they needed to show them how to work it out, what the resources were to help them and how to work with the competency framework.

> ‘They don’t care about definitions or the more theoretical aspects behind it all. But when we get to talk about EPAs and to show them the assessment tool…as long as I kept talking about EPAs, it didn’t mean anything to them. But, when I began writing EPAs for psychiatry…Oh, I get it!’ (PS-01/2014).

#### The Surgery Department

The Surgery Department Academic Leaders, recruited after 2012, spoke of their keen interest for pedagogy and their willingness to improve teaching in the department. One of them had been program director for many years (SU-02/2014), another in a previous life had been a physical educator and had completed her master’s degree in medical education (SU-01/2014). The Department Chair had known them since they were students and had already identified them as Academic Leaders. Their future in teaching:

> ‘must have shown clearly when we were students too, hum, that our…you know, even if you use the term ‘passion’, it’s…when you say it’s natural, and it already showed.’(SU-FG/2014)

Some mentioned that the recognition brought on by the official status of Academic Leader was part of what motivated them. Others mentioned that their decision to become Leaders was a natural consequence of their having chosen to work in a ‘university centre […] it’s part of my job, it’s something I like to do.’ (SU-FG/2014)

The first issue brought up by the Academic Leaders is the need to bring solutions to the teaching sites. This was seen as central to their work; their role is to help Faculty find new and alternative solutions to problems they encounter.

> ‘everybody sees that there are gaps here and there, but they don’t necessarily put in any requests [for assistance] because everyone does things the way they have been done traditionally since…you know, the same way…’(SU-01/2014)

Leaders in the Surgery Department also equated CBME implementation with an opportunity to improve teaching practice. Surgery Department Academic Leaders were aware from the outset they needed to work collectively on issues that didn’t coincide with their personal interests, but rather emanated from above, and were part of the general ‘top-down’ initiative to implement CBME. They said that other colleagues had refused the invitation to become Academic Leaders because they didn’t adhere to the project ‘it’s not true that they’ll make me talk about [pedagogy], only because they want me to talk about that.’ (SU-02/2014) This shift to a collective approach to CBME implementation marks the passage from the socialization stage to the externalization stage.

The externalization stage in surgery is also marked by resolution of issues of legitimacy. Some Leaders in surgery clearly had doubts about their ability to be Academic Leaders; ‘it’s not easy to be convinced that we can do it, that we are the right person that can necessarily do it because we took the week-long training.’ (SU-FG/2014) ‘What we need is that leaders be recognized in the different settings, especially in the programs, so that precisely, people come to us.’ (SU-FG/2014) At issue was the Academic Leader role, not in a position of formal authority, but a sort of expert consultant whose advice you can ignore if you wish:

> ‘So, the leader is something new, it’s not a program director, it’s not a division chief, it’s someone who will tell us that we have to teach more, in another way; who is going to give us work to do but we don’t know where he’s coming from and what title he has really. It wasn’t easy to deal with this.’ (SU-FG/2014).

In 2014, recently hired Academic Leaders realized that they needn’t bother explaining the CBME concepts to their colleagues, but they had to make them workable. They came to see that as their true mandate as Academic Leaders.

> ‘Well, people…sincerely….when we talk to people about EPAs, competencies, milestones, they have no idea what we’re talking about. And even if they attended presentations on the subject…I mean even myself who attended many of these presentations, it wasn’t at all clear at the beginning. So, I think people were wary at the beginning and I think they are still. And I understand, but I think that by doing it in practice it should be easier.’ (SU-FG/2014, 356-361)

The need to introduce EPAs for the assessment of competencies into surgery programs became clear to all instructors in the department and the Department Chair clearly expressed the need for it. This changed the nature of Leaders’ work because it wasn’t any longer a question of ‘training’ colleagues, but working with key people, such as Program Directors. Leaders consciously targeted mid-career attending surgeons in formal authority positions:

> ‘We turned to them because they were responsible for administrative tasks, for example, comorbidity and mortality or others. We went to see them and told them to do that because you already do it well in practice, so, do it for…and no one said no.’ (SU-FG/2014).

Surgery Department Leaders had clearly adopted the conception of their role as ‘behind the scenes’ advisor to persons in positions of authority. By making Program Directors present the EPAs during the pedagogy meetings, Academic Leaders managed to get them engaged in the process. However, they were acutely aware of their limited power:

> ‘I needed them to own the contents [of the EPAs] and that it would be their responsibility because as an Academic Leader, I couldn’t fathom the idea of investing all my energy to keep an eye on each surgeon in the ENT specialty program – and I think it needs to be like that. So, I just hope that [Program Directors] say “It’s my responsibility and I’ll do it”’(SU-FG/2014, 324-327)

The above quote and the following one illustrate how Leaders decided to take up a less visible role and to act as advisors to their colleagues. One of the Academic Leaders in the focus group said:

> ‘I relied heavily on the program director in ENT to start working as a leader. […] I told him that emails ‘have to come from you’. I could even write the emails for you and then you can send them on your behalf, this would help.’ (SU-FG/2014)

This Academic Leader expressed the hope that with time this would change, but that for the time being you had to partner up with a person in authority to push for CBME implementation. Another mark of the combination stage is the need for a yearly calendar of Department meetings devoted to CBME topics:

> ‘you know a sort of structure in which, we say there is such a number of meetings per year in such and such a context. You know, that’s it, there needed to be a sort of skeleton of a structure in which you can easily find yourself. This instead of constantly chasing each other to meet-up and work.’ (SU-FG/2014)

Finally, it needs to be said that Leaders were keenly aware that program accreditation requirements constituted a facilitating factor that made their work easier. They expressed it as the “big stick” to motivate people towards CBME implementation: ‘It is clear that accreditation was a great accelerator.’ (SU-FG/2014)

By 2014, Academic Leaders in the Surgery Department were becoming aware of what strategies would be most effective, especially that of working with Program Directors. The Department Chair had also provided support for Academic Leaders by instituting a calendar of meetings where information about pedagogy could be shared to all instructors in the Department.

#### The internalization stage in the three Departments

By 2014, Academic Leaders and Department Chairs in all three departments recognized that few people questioned the need to implement CBME anymore. Indeed, discourse on CBME was rarely cast in a negative light. Pedagogy became a recurring topic in most Department Continuing Medical Education meetings and yearly Pedagogy Days are offered by the departments explicitly focused on writing competency assessment tools. However, EPA implementation was just beginning and was far from becoming a widespread tool for teaching and assessment at the medical school.

## Discussion

This study presents a dynamic portrait of how the work of Academic Leaders for CBME implementation evolved over time. From an intuitive approach focused on ‘training’ colleagues on CBME, they adopt a collaborative approach, working within local academic committees and colleagues in positions of formal authority. This portrait affords unique insights into how Academic Leaders contributed to change.

At the socialization stage, CBME was an abstract concept and Academic Leaders simply relied on their intuition to ‘inform’ colleagues about it. As time passed, Academic Leaders realized that this approach was not the most effective. The realization that their target audience would not be reached this way, was key in their changing their approach. In order to gain traction among a greater audience, they reached out to create alliances. The sharing of their tacit knowledge, stemming from their questioning of assumptions, became crucial at this stage.

During the externalization stage, Academic Leaders came together with others who had received the same CBME training. This allowed them to acquire a common vocabulary, to share teaching materials and insights. CBME was becoming much more familiar to them and they felt more confident when explaining it to peers. Consequently, they were recognized as CBME experts within their Departments, and this consolidated their positions within the community and strengthened their resolve to persevere.

The transition from the externalization stage to the combination stage appears as a crucial step. Beside institutional supports (e.g. CAPHS logistical support, time off and explicit vision from Chairs expressed in Department meetings), Leaders’ work was clearly facilitated by the introduction of EPAs. Academic Leaders saw the value of this tool for CBME implementation and became key contributors for their development with colleagues. Another crucial step ensued: Academic Leaders worked together with Program Directors or Pedagogic Committees to develop Department specific EPAs with peers. At this stage, Leaders perceived that CBME was becoming the norm and that resistance to it was diminishing.

It is undeniable that CBME implementation was facilitated in large part by clear institutional support, the appearance of EPAs and the fact that CBME was required by the national accreditation agency. What our study focused on is how Academic Leaders carried this process within their communities. Their work in this context can best be summarized by : collaborating among each other to strengthen their mastery of abstract notions inherent in collective project, securing recognition as experts from peers, transitioning from a role of ‘preacher’ to one of ‘advisor’, and actively participate in the writing of tools and teaching materials within departmental committees.

In sum, by questioning current teaching practice and engaging with peers at first and then with Faculty members in positions of authority and committees, Academic Leaders figured out how to persuade their peers to adopt CBME. These results provide evidence that the work of Academic Leaders is to translate vision statements and goals adopted by institutional authorities into workable pieces of information that make practical sense to their peers.

### Study limitations

In order to examine the evolution of the Leader’s role and to test the use of the organizational knowledge creation model we selected the case study approach with embedded levels. This meant that the individual differences amongst individual leaders themselves were not taken into account. The cost is that factors such as personality, medical specialties, hospital settings, etc. that could play a role were overlooked. These factors will have to be the focus of further research on the topic.

However, focusing expressly on the relationships between leaders and the social structures in which they act is of sufficient value to offset the cost. The impact of these relationships on organizational change cannot be underestimated and their examination can provide valuable insight on how Academic Leaders contribute to it. Also, our selection of the diverse sampling method allows us to be confident that the results constitute a valid and credible portrayal of the evolution of Academic Leaders’ work in the implementation of competency-based education in institutions of higher learning.

## Conclusion

Implementing CBME is considered an important organizational change for medical schools. Getting Faculty to envision such change is a daunting challenge. Given the nature of academia, in which Faculty enjoy academic freedom, the challenge appears unachievable, even if improving teaching practice becomes a requirement imposed by accreditation agencies.

Academic Leadership is a concept that has been introduced as a strategy to overcome these challenges. This study illustrates one such example where Academic Leaders were trained and supported to convince their peers that change was necessary and possible. In order to map these changes, the organizational knowledge creation framework yielded descriptions of stages in the evolution of the work over two years that allowed us to identify specific factors that facilitated change. We hope that Higher Education Institutions authorities and researchers engaged in reforming their organizations will find it useful to use this framework to track progress towards change.

## References

1. Aquilante AG, da Silva RF, de Souza MBB, Kishi RGB. Implementation of Competency-Based Curriculum in Medical Education: Perspective of Different Roles. ISRN Education. 2012;2012:7.

2. Frank JR, Mungroo R, Ahmad Y, Wang M, De Rossi S, Horsley T. Toward a definition of competency-based education in medicine: a systematic review of published definitions. Medical Teacher. 2010;32(8):631–7.

3. Frank JR, Snell LS, ten Cate O, Holmboe ES, Carraccio C, Swing SR, et al. Competency-based medical education: theory to practice. Medical Teacher. 2010;32(8):638–45.

4. Schultz K, McEwen L, Griffiths J. Applying Kolb’s Learning Cycle to Competency-Based Residency Education. Acad Med. 2016;91.

5. Schultz K, Griffiths J. Implementing Competency-Based Medical Education in a Postgraduate Family Medicine Residency Training Program: A Stepwise Approach, Facilitating Factors, and Processes or Steps That Would Have Been Helpful. Academic Medicine. 2016;91(5):685–9.

6. McEwen LA, Griffiths J, Schultz K. Developing and Successfully Implementing a Competency-Based Portfolio Assessment System in a Postgraduate Family Medicine Resideny Program. Academic Medicine. 2015;90(11):1515–26.

7. Whitehead C, Kuper A. Competency-based training for physicians: Are we doing no harm?. Canadian Medical Association Journal. 2014;187(4):E128–E9.

8. Delener N. Leadership excellence in higher education: Present and future. Contemporary Issues in business and government. 2013;19(1):19–33.

9. Carless SA, Wearing, A.J., Mann, L.. A Short Measure of Transformational Leadership. Journal of Business and Psychology. 2000;14(3):389–405.

10. Kezar A, Carducci R, Contreras-McGavin M. Rethinking the “L>” Word in Higher Education: The Revolution of Research on Leadership: ASHE Higher Education Report, Volume 31, Number 6. K. Ward LW-W, editor. Hoboken, NJ: Josey-Bass; 2006. 240 p.

11. Kezar A, Lester J. Supporting Faculty Grassroots Leadership. Research in Higher Education. 2009;50:715–40.

12. Abbasi E. The role of transformational leadership, organizational culture and organizational learning in improving the performance of Iranian agricultural faculties. High Educ. 2013;66:505–19.

13. Wisniewski MA. Leadership and higher education: Implications for leadership development programs. Academic Leadership Journal. 2004;2.

14. Juntrasook A. ‘You do not have to be the boss to be a leader’: contested meanings of leadership in higher education. Higher Education Research & Development. 2014;33(1):19–31.

15. House R, Javidan M, Dorfman P. Project GLOBE: An introduction. Applied Psychology: An International Review. 2001;50(4):489–505.

16. Leaming DR. Academic Leadership: A Practical Guide to Chairing the Department. San Francisco: Jossey-Bass; 2007.

17. Ramsden P. Learning to Lead in Higher Education. London: Routledge; 1998.

18. Albanese MA, Mejicano, G., Anderson, W.M. & Gruppen, L.. Building a competency-based curriculum: the agony and the ecstasy. Advances in Health Sciences Education. 2010;15(3):439–54.

19. French WL, Bell C, Zawacki RA. Organization development and transformation: Managing effective change. 6th Special Indian Edition ed. New Delhi: Tata McGraw-Hill Education Private Ltd.; 2006.

20. Krupat E, Pololi L, Schnell ER, Kern DE. Changing the culture of academic medicine: the C-Change learning action network and its impact at participating medical schools. Academic Medicine. 2013;88(9):1252–8.

21. Giddens A. The Constitution of Society: Outline of the Theory of Structuration. Berkeley, CA: University of California Presse; 1984.

22. Ashforth BE, Mael, F.. Social Identity and the Organization. The Academy of Management Review. 1989;14(1):20–39.

23. Stensaker B. Organizational identity as a concept for understanding university dynamics. High Educ. 2015;69:103–15.

24. Wenger E. Communities of Practice: Learning, Meaning and Identity. Cambridge, UK: Cambridge University Press; 1998.

25. Billett S. Guided learning at work. Journal of Workplace Learning. 2000;12(7):272–85.

26. Baumgard P. Tacit Knowledge in Organizations. London: Sage Publications; 1999.

27. Polanyi M. Personal Knowledge: Towards a Post-Critical Philosophy. Chicagi, IL: The University of Chicago Press; 1968/ 2015.

28. Polanyi M. The Tacit Dimension 1966.

29. Nonaka I, Tekeuchi H. The Knowledge-Creating Company: How Japanese Companies Create the Dynamics of Innovation. Oxford: Oxford University Press; 1995.

30. Nonaka Ik, Toyama R, Konno N. SECI, Ba and Leadership: A Unified Model of Dynamic Knowledge Creation. Long Range Planning. 2000;33:5–34.

31. Nonaka I, Toyama, R. The knowledge-creating theory revisited: knowledge creation as a synthesizing process. Knowledge Management Research & Practice. 2003;1:2–10.

32. Buckley PF. The medical school dean: leadership and workforce development. Acad Psychiatry. 2014;38(1):82–5.

33. Jones S, Lefoe G, Harvey M, Ryland K. Distributed leadership: A collaborative framework for academics, executives and professionals in higher education. Journal of Higher Education Policy and Management. 2012;34(1):67–78.

34. Yin RK. Case Study Research : Design and Methods (5th edition). Thousand Oaks, CA: Sage Publications; 2014.

35. Gerring J. Techniques for Choosing Cases. In: Gerring J, editor. Case study research: principles and practices. Cambridge: Cambridge University Press; 2007. p. 86–150.

36. Boyatzis RE. Transforming qualitative information: Thematic analysis and code development. Thousand Oaks, CA: Sage Publications; 1998.

37. Cresswell JW. Research Design: Qualitative, Quantitative, and Mixed Methods Approaches, 2nd edition. Thousand Oaks, CA: Sage Publications, Inc.; 2003. 246 p.

38. Peters H, Holzhausen Y, Boscardin C, ten Cate O, Chen HC. Twelve tips for the implementation of EPAs for assessment and entrustment decisions. Medical Teacher. 2017;Online.

39. ten Cate O, Scheele, F. Competency-Based Postgraduate Training: Can We Bridge the GAP between Theory and Clinical Practice? Academic Medicine. 2007;82(6):542–7.

